# Disconnect Between Succinate, Succinyl-CoA, and Histone Succinylation in Cells and Gastrointestinal Stromal Tumors with Succinate Dehydrogenase Deficiency

**DOI:** 10.64898/2026.10.05.756882

**Authors:** Adam Chatoff, Trisha Gullapudi, Sun Venkat, Talha Chaudhry, Mariola Marcinkiewicz, Jane Koshy, Kiara Hackey, Kathy Cai, Sarah Graff, Stephanie Stransky, Simone Sidoli, Lori Rink, Nathaniel W. Snyder

## Abstract

**Background:** Succinate dehydrogenase (SDH) is a heterotetrameric enzyme in both the tricarboxylic acid (TCA) cycle and the electron transport chain. SDH-deficiency has been shown to drive the oncogenesis of cancer types including gastrointestinal stromal tumors (GIST), paraganglioma/pheochromocytoma, and clear cell renal cell carcinoma. Previous work on SDH loss has shown increases in succinate, decreases in fumarate, as well as increase in the succinate to α-ketoglutarate ratio, and correlated these changes to metabolite signaling and epigenetic dysregulation. Alterations to other TCA intermediates, including succinyl-CoA, the acyl-donor for succinylation, and the resulting cellular effects has not been fully explored.

**Methods:** We used liquid chromatography high resolution mass spectrometry to quantify TCA cycle metabolites across the NCI-60 panel of cell lines, three SDH-competent and SDH-deficient model systems, and human tumor samples. We measured histone post-translational modifications in SHD-deficient and SDH-competent models and examined the correlation and potential pharmacological targeting of described desuccinylase enzymes by protein and cell assays.

**Results:** We found an increase in succinyl-CoA in multiple SDH-deficient models, but no correlation between succinate to fumarate ratio and succinyl-CoA concentrations in the NCI-60. Interestingly, we found that while there was an increase in succinyl-CoA in the SDH-deficient cells, there was no corresponding increase in histone succinylation. We also observed a decrease in lysine malonylation and glutarylation, two other anionic lysine modifications. Cell viability and clonogenic assays revealed very weak or no selective effect of desuccinylase targeting in SDH-deficient models. Similarly, using SDH-deficient versus SDH-competent GIST patient samples, we examined both protein expression and metabolite abundance at the bulk tumor level and found no significant increase in succinyl-CoA or desuccinylase expression, despite the increase in succinate to fumarate ratio.

**Conclusions:** We found that SDH-deficiency led to altered central carbon metabolism including increases to succinate and succinyl-CoA. Despite data that suggested the upregulation of desuccinylases may mediate a disconnect between succinyl-CoA concentration and histone succinylation, we found weak or no reductions in cell viability in response to pharmacological inhibition of desuccinylases. Additionally, no correlation of SDH loss with desuccinylase expression and succinyl-CoA in bulk tumor samples was observed.

## BACKGROUND

The tricarboxylic acid cycle (TCA) and electron transport chain (ETC) are central metabolic pathways that generate energy and biosynthetic precursors for cell proliferation and survival [1]. Succinate dehydrogenase is a heterotetrameric enzyme embedded in the inner mitochondrial membrane and is the only enzyme to play a role in both the TCA and ETC. SDH catalyzes the conversion of succinate to fumarate in the forward direction within the TCA cycle which is coupled to the reduction of ubiquinone to ubiquinol feeding electrons to the ETC [2]. Loss of SDH expression has been shown to drive a variety of cancer types, including gastrointestinal stromal tumors (GIST), pulmonary chondromas, paragangliomas/pheochromocytomas, and clear cell renal cell carcinoma [3-5]. Recently, it has been reported that a functional SDH-deficiency is a common feature of clear cell renal cell carcinoma, even when driven by a different oncogenic mutation [6]. Additionally, SDH-deficiency has been shown to cause Leigh Syndrome, a neurodegenerative condition [7].

Loss of SDH expression leads to metabolic alterations. Prior work has shown that SDH-deficiency leads to the accumulation of succinate and a reduction in fumarate [8, 9]. Additionally, SDH-deficiency leads to an increased succinate to α-ketoglutarate ratio [9]. This increase of metabolite ratio leads to product inhibition of α-ketoglutarate dependent dioxygenases including DNA and histone demethylases and proline hydroxylase, which mark hypoxia inducible factor 1 for degradation [10-12]. This has led existing studies in the epigenetic landscape of SDH-deficiency to primarily focus on methylation status.

Despite the central position of SDH within the ETC and the TCA cycle, alterations to other TCA intermediates have not been previously explored. Specifically, succinyl-CoA, one step removed from the SDH catalyzed reaction in the reductive TCA, is a reactive metabolite with established functions beyond serving as an intermediate in the steps between α-ketoglutarate and succinate. Succinyl-CoA is the donor for lysine succinylation, an anionic post-translational modification that alters protein stability, activity, and subcellular localization [13]. Histone lysine succinylation is thought to be positively correlated with succinyl-CoA [14] and alters gene expression due to histone lysine succinylation [15].

To investigate alterations to succinyl-CoA concentrations and succinylation status we made use of 2 isogenic cell models, disease relevant cell models, as well as patient samples. To better understand the generalizability of SDH-loss in common cancer cell lines, we also performed targeted metabolomics on the NCI-60 panel. Lastly, we made use of SDH-competent and SDH-deficient patient samples from the Fox Chase Cancer Center Biorepository. This diversity of models allows these studies to investigate the SDH-deficiency driven alterations to succinylation status in disease relevant context. Given that SDH-deficiency can drive multiple pathologies, our findings will be translatable to other SDH-deficient contexts.

## METHODS

### Cell culture and cell lines

143B cell lines were cultured in complete DMEM (Gibco), high glucose medium supplemented with 10% fetal bovine serum (Gemini), 1% penicillin/streptomycin (Gibco), 1mM sodium pyruvate (Gibco) and 0.1mg/mL uridine (Sigma). Hap1 cell lines were cultured in complete IMDM (Gibco) supplemented with 10% fetal bovine serum (Gibco) and 1% penicillin/streptomycin (Gibco). The human 143B osteosarcoma cells were generated as previously described [8] and was gifted from the laboratory of Dr. Jessica Spinelli. The human HAP1 leukemia cell lines were generated as previously described [16] and were gifted from Dr. Michael Heinrich. All cells were grown in a humidified atmosphere at 37°C containing 5% CO_2_ and were routinely tested for mycoplasma through PCR analysis using specific primers (fwd: ACACCATGGGAGYTGGTAAT and rev: CTTCWTCGACTTYCAGACCCAAGGCAT).

### Colony formation assay

The class 1 HDAC inhibitor Trichostatin A was purchased from Sigma. 250 cells were plated in triplicates in 6-well plates in triplicates and incubated in media containing 40 nM, 10 nM, 2.5 nM TSA or DMSO only for 12 days with treatment media changing every 3 days. At the end of the experiment, colonies were stained with 0.5% crystal violet (Sigma) diluted in 20% methanol in water. Following incubation for 20 min with gentle agitation at RT, cells were washed for 3 times with water and air dried overnight. Colony number and colony size was quantified using ImageJ.

### Metabolite extraction and LC-MS

For extraction of acyl-CoAs, culture medium was aspirated from cells in 10-cm plates before adding 1 ml of ice-cold 10% trichloroacetic acid to plates. Plates were scraped on ice to collect cells. Internal standard was added to each sample containing [^13^C_3_^15^N_1_]-labeled acyl-CoAs generated in pan6-deficient yeast culture [17]. Samples were sonicated for 10 x 0.5 s pulses and centrifuges at 16,000g for 10 min at 4 °C. The supernatant was collected and purified by solid-phase extraction using Oasis HLB 1cc (30 mg) solid phase extraction (SPE) columns (Waters). Eluate was dried down under nitrogen gas and samples were resuspended in 50μl of 5% 5-sulfosalicylic acid (w/v) and transferred to a 96-well plate. Samples were analyzed by an Vanquish Duo HPLC (Thermofisher) coupled to a Thermo QExactive Plus instrument in positive electrospray ionization mode as previously described [18]. For quantitation, a calibration curve was generated using commercially available unlabeled standards (Sigma) For extraction of tricarboxylic acid cycle intermediates, cells were washed three times with 1ml ice-cold 0.9% NaCl to remove extracellular metabolites and scraped on ice in 1ml -80°C 80% Optima grade methanol/20% Optima grade water (Thermo). Extracts were collected in 1.5ml tubes and 50μl of 1μM of 13C TCA Cycle internal standard (MedChemExpress) was added to each sample. Samples were vortexed and incubated at -80°C for 30 min following centrifugation at 18,000g for 10 min at 4°C. Supernatants were transferred into a 96-well plate and dried under nitrogen gas at room temperature overnight. Dried metabolites were resuspended in 50μl 95% Optima grade water/ 5% Optima grade methanol (Thermo). For each sample, 2μl was injected and analyzed using a Vanquish Duo UHPLC system coupled to a Thermo QExactive Plus Orbitrap Instrument in ion switching electrospray ionization mode in full scan mode from 70-1000 m/z. The HPLC system used a hydrophilic interaction chromatography (HILIC) analytical column (SeQuant ZIC-pHILIC). The column was kept at 25 °C and the flow rate was 0.15ml/min. The mobile phase was solvent A (25 mM ammonium carbonate and 0.1% ammonia hydroxide in water) and solvent B (Optima Acetonitrile). Elution gradients were run starting from 80% B to 20% B from 0-20 min; 80% B was held from 20-28 min. For each analyte and the internal standard, the peak corresponding to the [M+H]^+^ or [M-H]^-^ ion at 5ppm was integrated in Tracefinder 4.1 (Thermo Scientific).

### Histone extraction and digestion

Histone proteins were extracted from the pellet as described by [19, 20] to ensure good-quality identification and quantification of single histone marks. Briefly, histones were acid-extracted with chilled 0.2 M sulfuric acid (5:1, sulfuric acid: pellet) and incubated with constant rotation for 4 h at 4°C, followed by precipitation with 33% trichloroacetic acid (TCA) overnight at 4°C. Then, the supernatant was removed, and the tubes were rinsed with ice-cold acetone containing 0.1% HCl, centrifuged and rinsed again using 100% ice-cold acetone. After the final centrifugation, the supernatant was discarded and the pellet was dried using a vacuum centrifuge. The pellet was dissolved in 50 mM ammonium bicarbonate, pH 8.0, and histones were subjected to derivatization using 5 µL of d_10_-propionic anhydride (Sigma Aldrich) which allows analysis of endogenous propionylation, and 14 µL of ammonium hydroxide (Sigma Aldrich) to balance the pH at 8.0. The mixture was incubated for 15 min and the procedure was repeated. Histones were then digested with 1 µg of sequencing grade trypsin (Promega) diluted in 50mM ammonium bicarbonate (1:20, enzyme:sample) overnight at room temperature. Derivatization reaction was repeated to derivatize peptide N-termini. The samples were dried in a vacuum centrifuge.

### Sample desalting

Prior to mass spectrometry analysis, samples were desalted using a 96-well plate filter (Orochem) packed with 1 mg of Oasis HLB C-18 resin (Waters). Briefly, the samples were resuspended in 100 µl of 0.1% TFA and loaded onto the HLB resin, which was previously equilibrated using 100 µl of the same buffer. After washing with 100 µl of 0.1% TFA, the samples were eluted with a buffer containing 70 µl of 60% acetonitrile and 0.1% TFA and then dried in a vacuum centrifuge.

### Histone PTM LC-MS/MS Acquisition and Analysis

Samples were resuspended in 10 µl of 0.1% TFA and loaded onto a Dionex RSLC Ultimate 300 (Thermo Scientific), coupled online with an Orbitrap Fusion Lumos (Thermo Scientific). Chromatographic separation was performed with a two-column system, consisting of a C-18 trap cartridge (300 µm ID, 5 mm length) and a picofrit analytical column (75 µm ID, 25 cm length) packed in-house with reversed-phase Repro-Sil Pur C18-AQ 3 µm resin. Peptides were separated using a 30 min gradient from 1-30% buffer B (buffer A: 0.1% formic acid, buffer B: 80% acetonitrile + 0.1% formic acid) at a flow rate of 300 nl/min. The mass spectrometer was set to acquire spectra in a data-independent acquisition (DIA) mode. Briefly, the full MS scan was set to 300-1100 m/z in the orbitrap with a resolution of 120,000 (at 200 m/z) and an AGC target of 5°10e5. MS/MS was performed in the orbitrap with sequential isolation windows of 50 m/z with an AGC target of 2°10e5 and an HCD collision energy of 30.

Histone peptides raw files were imported into EpiProfile 2.0 software [21]. From the extracted ion chromatogram, the area under the curve was obtained and used to estimate the abundance of each peptide. In order to achieve the relative abundance of post-translational modifications (PTMs), the sum of all different modified forms of a histone peptide was considered as 100% and the area of the particular peptide was divided by the total area for that histone peptide in all of its modified forms. The relative ratio of two isobaric forms was estimated by averaging the ratio for each fragment ion with different mass between the two species. The resulting peptide lists generated by EpiProfile were exported to Microsoft Excel and further processed for a detailed analysis.

### Protein extraction and western blotting

For protein isolation, cell pellets were washed with 1X PBS and resuspended in lysis buffer (50 mM Tris-HCL pH 8.0, 2 mM EDTA, 100 mM NaCL, 1% Triton-X-100, 10% glycerol, 0.5 mM PMSF and 1X protease inhibitor cocktail). Samples were sonicated for 5 x 0.5s pulses and lysates were centrifuged at 10,000 rpm for 10 min at 4°C. Total protein concentration was quantified by BCA assay (Thermo Scientific). Thirty micrograms of protein extracts were diluted in 4X laemmli sample buffer (Bio-Rad) containing DTT (Sigma) and denatured at 95 °C for 5 min. Proteins were separated on 4-12% gradient SDS-PAGE gels (BioRad) and transferred to nitrocellulose membranes (Bio-Rad). The blots were stained with Ponceau S Staining solution (Cell Signaling) and following blocking with 5% BSA in PBST for 1 h at RT, membranes were incubated with primary antibodies in 5% BSA in PBST overnight at 4°C with agitation. Membranes were then incubated with fluorescently labelled secondary antibodies diluted in 5% BSA in PBST for 1 h at RT. All membranes were developed using a LI-COR Odyssey CLx system. Antibodies used in this study were as follows: SDHA, -B, -C(Abcam), HDAC1, 2, 3 (Abcam), SIRT5, 7 (Abcam), α-Tubulin (Cell Signaling).

### Slide Preparation and Immunohistochemistry

Immunohistochemical (IHC) staining for SDHB, SIRT5/7, HDAC1/2/3 was performed on formalin-fixed, paraffin-embedded (FFPE) tissue sections cut at 5 µm using an automated staining platform (Roche Ventana Discovery Ultra). Tissue sections were dried overnight at 50 °C prior to staining.

Slides were deparaffinized onboard using EZ Prep solution according to the manufacturer’s protocol. Heat-induced epitope retrieval (HIER) was performed using Cell Conditioning 1 (CC1; Tris-EDTA buffer, pH 8.0) at 95 °C for 32 minutes. Retrieval conditions were optimized during assay development to achieve optimal signal-to-noise ratios for antibody detection.

Primary antibody incubation was carried out at room temperature under the following conditions; Anti-SDHB antibody (Abcam, ab14714), dilution 1:100, incubation for 60 minutes; Anti-SIRT5 antibody (Abcam, ab259967), dilution 1:100, incubation for 60 minutes; Anti-SIRT7 antibody (Abcam, ab259968), dilution 1:50, incubation for 60 minutes; Anti-HDAC1 antibody (Abcam, ab109411), dilution 1:200, incubation for 32 minutes; Anti-HDAC2 antibody (Abcam, ab32117), dilution 1:800, incubation for 32 minutes; Anti-HDAC3 antibody (Abcam, ab ab32369), dilution 1:200, incubation for 32 minutes. Detection was performed using a multimer-based horseradish peroxidase (HRP) system (Discovery OmniMap Anti-Rabbit HRP). Signal visualization was achieved using the Discovery ChromoMap DAB IHC Detection Kit, with 3,3′-diaminobenzidine (DAB) serving as the chromogenic substrate to produce a brown precipitate at antigen sites. Slides were subsequently counterstained with hematoxylin followed by application of a bluing reagent. Negative controls were generated by substituting the primary antibody with species-matched non-immune IgG.

### Slide Scanning

All stained sections were digitized as whole-slide images (WSIs) using a Hamamatsu NanoZoomer S60 slide scanner at 40× magnification under standard acquisition settings.

### Image Analysis Using Visiopharm

Quantification of SDHB, SIRT6/7, HDAC1/2/3 expression was performed using customized analysis applications developed in Visiopharm software (Visiopharm A/S, Denmark; version 2025.08, 64-bit). All applications were optimized to accommodate biomarker-specific staining characteristics and ensure reproducible measurements across samples.

### Tumor Detection

Tumor detection applications were developed to distinguish viable tumor regions from surrounding stromal and necrotic tissue. Quantitative analysis was restricted to automatically detected tumor areas to ensure consistent exclusion of non-tumor tissue. The detection algorithms were customized to accommodate morphological variability across samples, enabling consistent tumor identification while minimizing inclusion of background or nonviable tissue.

### H-Score Application Development

Following tumor region detection, color deconvolution was applied to separate hematoxylin and DAB channels. DAB-positive pixels were classified into high-, medium-, and low-intensity categories, while hematoxylin-positive pixels were detected for normalization. Analysis parameters were optimized for stain intensity, cellular morphology, and background characteristics unique to each antibody. Threshold levels, color deconvolution settings, and tissue classification criteria were iteratively refined to achieve consistent and biologically meaningful segmentation across all samples. Threshold values for both channels were optimized independently for SIRT5/7, HDAC1/2/3 to reflect biomarker-specific staining patterns. Areas corresponding to high-, medium-, and low-intensity DAB staining, along with hematoxylin-positive areas, were calculated. Cytoplasmic pixel-based H-scores (Cyto H-scores) were computed using the formula: pix H-score = (3 × H_p_ + 2 × M_p_ + 1 × L_p_) / N_p_ where H_p_, M_p_, and L_p_ represent areas of high-, medium-, and low-intensity DAB-positive pixels, respectively, and N_p_ represents the hematoxylin-positive area. Sequential analysis prioritized the DAB channel to ensure accurate cytoplasmic quantification while minimizing interference from nuclear staining.

## RESULTS

### SDH-Deficiency Alters Tricarboxylic Acid Cycle Abundance Including Increases in Succinate and Succinate to Fumarate Ratio

To study how SDH-deficiency alters TCA intermediate abundance in our model systems, we utilized stable isotope dilution liquid chromatography-high resolution mass spectrometry (LC-HRMS) to quantify polar metabolites including succinate, fumarate, and α-ketoglutarate. Since cell volume may be different across lines, we used Coulter principle-based counting to measure cell volume. We found that in our 143B model systems, SDH-deficiency increased succinate from μM cellular concentrations (WT 129.0 μM mean, 14.81 μM std. dev.) to mM (SDHB KO 1,871 μM mean, 79.37 μM std. dev.), increased succinate/ fumarate ratio and the succinate/α-ketoglutarate ratio (**Figure 1A-1B**) consistent with previous studies on SDH-deficiency [8, 9]. The Hap1 model system recapitulated these findings with SDH-deficiency significantly increased succinate concentrations (WT 63.39 μM mean, 11.35 μM std. dev., SDHA KO 858.3 μM mean, 71.01 μM std. dev., SDHA R31x 588 μM mean, 67.52 μM std. dev.), the succinate/fumarate ratio and the succinate/α-ketoglutarate ratio. SDH-loss in 143B and Hap1 models produced very similar succinate/fumarate ratios, and although succinate/α-ketoglutarate ratio increase was observed in both models, the absolute value of the increase was markedly different. We next quantified polar metabolites in SDH-competent and SDH-deficient gastrointestinal stromal tumor (GIST) patient tissues flash frozen after surgical resection. We again found that SDH-deficient bulk GIST tissues had increased succinate (SDH-competent GIST 58.32 nmol mean, SDH-deficient GIST 1580 nmol mean) and succinate/α-ketoglutarate ratio (**Figure 1C**). Although the median of the succinate/fumarate ratio for SDH-deficient samples, with some tumor samples reaching ratios of over 100, was substantially higher than the SDH-competent controls, the difference was not statistically significant, which suggests that tumor heterogeneity complicates this analysis, especially in the context of a rare cancer where accrual of additional samples may be logistically difficult (**Figure 1G**).

**Figure 1:**
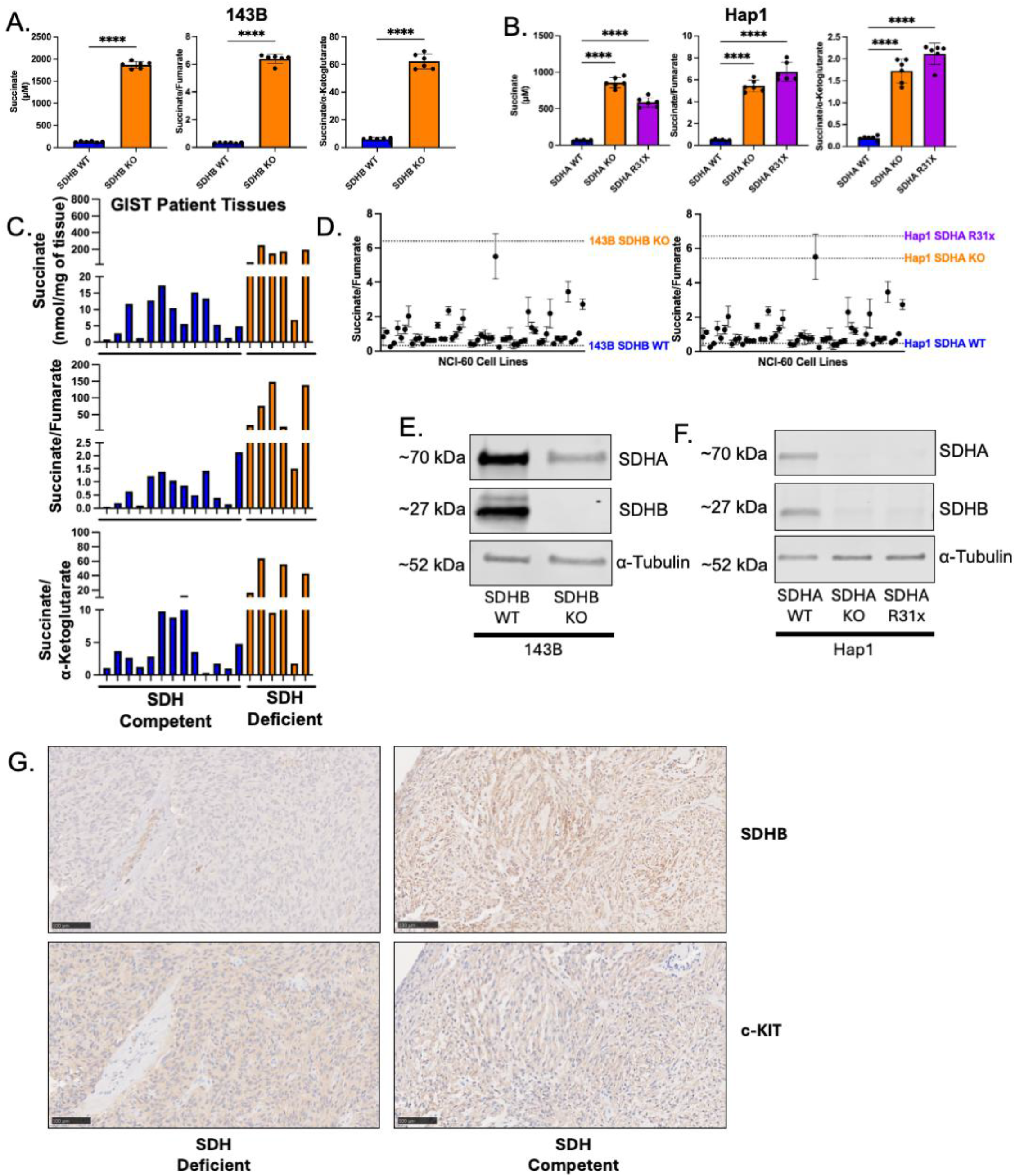
SDH-Deficiency Alters Tricarboxylic Acid Cycle Abundance Including Increases in Succinate and Succinate to Fumarate Ratio. **A**. Succinate concentration and succinate/fumarate ratio in 143B SDHB WT and SDHB KO (n = 6). **B**. Succinate concentration and succinate/fumarate ratio in Hap1 SDHA WT, SDHA KO, SDHA R31x (n = 6). **C**. Succinate concentration and succinate/fumarate ratio in SDH-competent (n = 12) and SDH-deficient GIST patient samples (n = 6). **D**. Succinate to fumarate ratio in NCI-60 cell lines (n = 3). Lines mark 143B SDHB WT and KO and Hap1 SDHA WT, KO and R31x Succinate/Fumarate Ratios. **E**. Western blot analysis of SDHA and SDHB in 143B SDHB WT and SDHB KO with alpha-Tubulin as loading control. **F**. Western blot analysis of SDHA and SDHB in Hap1 SDHA WT, SDHA KO, and SDHA R31x with alpha-Tubulin as loading control. **G**. Representative SDHB and c-KIT IHC images in SDH-competent and SDH-deficient GIST patient tissues. Significance determined by two-tailed T-Test for 143B and GIST Tissue Analyses. Significance determined by one-way ANOVA for Hap1 Analyses * p < 0.05 *** p < 0.001 **** p < 0.0001.

Given the wide range in succinate/fumarate ratio between our Hap1, 143B cell lines, and the SDH-deficient GIST cell lines, we systematically investigated the succinate/fumarate ratio across the NCI-60 panel of cell lines [22, 23]. Lines were validated by short tandem repeat profiling and cell line authentication before use. We extracted polar metabolites from 57 of the 60 cell lines and quantified the succinate/fumarate ratio. We found that only one of the NCI-60, H226, a male human lung epithelial squamous cell carcinoma or mesothelioma derived line, had a succinate/fumarate ratio similar to the Hap1 and 143B SDH knockout cell models (**Figure 1D, 1E, 1G)**. To our knowledge, there are no reports in the literature demonstrating that H226 is SDH-deficient.

### SDH-Deficiency Alters Acyl-CoA Abundance in 143B and Hap1 Cell Models, but Not in Bulk Human Patient Samples

Previous work in describing the metabolome in SDH-deficiency has focused on alterations to succinate, fumarate, and α-ketoglutarate concentrations. However, other TCA metabolites have been largely not reported, including acyl-CoAs. We hypothesized that SDH-deficiency would lead to an accumulation of succinyl-CoA, only one step removed from the SDH-catalyzed step in the TCA cycle. To investigate this, we profiled acyl-CoA alterations by LC-HRMS. We found that in the 143B and Hap1 model systems, SDH-deficiency resulted in an accumulation of succinyl-CoA, but not other CoAs including propionyl-CoA (**Figure 2A-2B**). Acetyl-CoA was inconsistently altered, with an increase in the 143B SDHB KO, but decreased in Hap1 SDHA KO (**Figure 2A-2B)**. We then tested the acyl-CoA profile of SDH-competent and SDH-deficient GIST patient samples. Despite the previously observed accumulation of succinate in the SDH-deficient samples, we saw no increase in succinyl-CoA in our SDH-deficient tissues, nor an increase in succinyl-CoA/CoASH ratio (**Figure 2C**). While this is inconsistent with our initial hypothesis and findings in our 143B and Hap1 model systems, the SDH-deficient acyl-CoA profile may be masked by SDH-competent stromal and immune cells found in the tumor microenvironment. SDH-deficient GIST are characterized as being well vascularized [24-26], so there may be significant SDH-competent cell infiltration. Additionally, unlike TCA metabolites including succinate, acyl-CoAs are not freely excreted into the extracellular environment [27]. As such, excess succinyl-CoA cannot build up in the tumor microenvironment as is possible for succinate.

**Figure 2:**
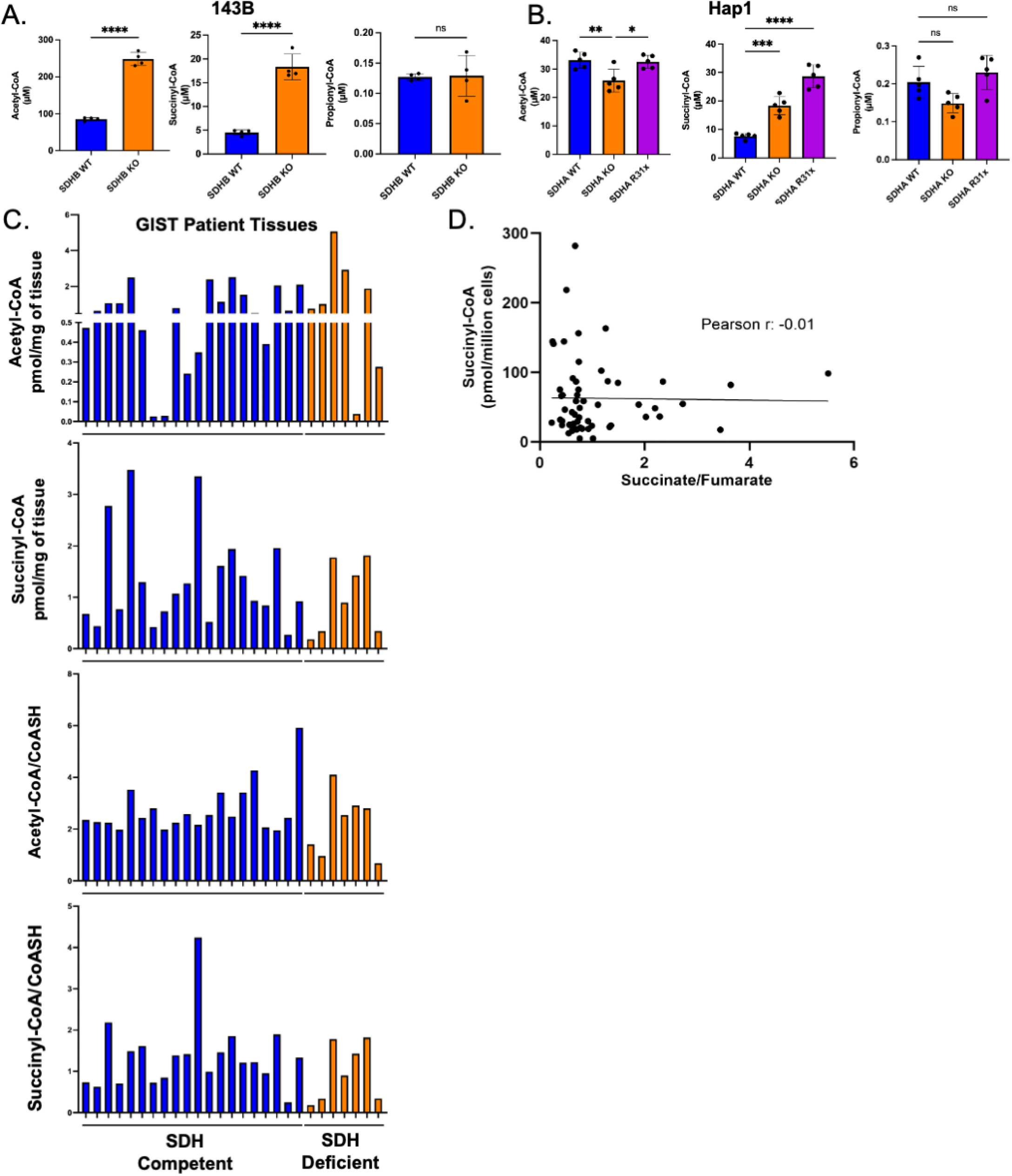
SDH-Deficiency Alters Acyl-CoA Abundance in 143B and Hap1 Cell Models, but Not in Bulk Human Patient Samples. **A**. Acyl-CoA quantification in 143B SDHB WT and SDHB KO cells (n = 4). **B**. Acyl-CoA quantification in Hap1 SDHA WT, SDHA KO, and SDHA R31x cell lines (n = 5). **C**. Acyl-CoA Quantification in SDH-Competent (n = 12) and SDH-Deficient GIST Patient Samples (n = 7). **D**. Correlation between Succinate/Fumarate ratio and Succinyl-CoA concentration. Significance determined by two-tailed T-Test for 143B and GIST Tissue Analyses. Significance determined by one-way ANOVA for Hap1 and GIST Cell Line Analyses. Pearson’s r value calculated by Pearson’s Correlation test. * p < 0.05 *** p < 0.001 **** p < 0.0001.

We also investigated if succinate/fumarate ratio was correlated with amount of succinyl-CoA. We quantified the acyl-CoA pool in the NCI-60 cell lines and found that there was no correlation between succinate/fumarate ratio and succinyl-CoA in these lines (**Figure 2D)**. This may indicate that SDH-deficiency leads to succinyl-CoA concentration changes through alternative mechanisms than accumulation of succinate. Another possible explanation is that SDH-competent cell lines do not accumulate enough succinate to alter the succinyl-CoA pool.

### SDH-Deficiency Alters Histone PTM Status Including Decreases in Desuccinylase Sensitive Marks

Succinylation is a post-translational modification where the succinyl group of succinyl-CoA is transferred to the free primary amine group of a lysine. Succinylation is an anionic modification at cellular pH, thus, switches the lysine positive charge to a negative charge and can alter the subcellular localization, function and stability of a protein [13]. Histones can be succinylated and previous work has shown that histone succinylation can alter chromatin structure and gene expression [28-31]. Given the observed increase in succinyl-CoA in our SDH-deficient 143B and Hap1 models, we sought to identify if there was a corresponding increase in histone succinylation. To investigate this, we made use of a previously described mass spectrometry-based bottom-up proteomics method to quantify alterations to the histone post-translational (PTM) landscape [20]. We extracted histones from our 143B, Hap1, and GIST patient samples.

Interestingly, and contrary to our initial hypothesis, we found that there was no significant increase in succinylation in our SDH-deficient 143B cell lines (**Figure 3A**). In our Hap1 models, we only saw a single increased succinylation site, H4K31, in the SDHA KO lines (**Figure 3B)**. In the SDHA R31x lines, we saw no increase in succinylation (**Figure 3C)**. In our SDH-deficient patient samples, in fact, we saw a decrease in multiple succinylation sites (**Figure 3D**). These findings in the Hap1 and 143B models were surprising given that we saw an increase in succinyl-CoA in our SDH-deficient 143B and Hap1 model systems. In addition, in our 143B models SDH-deficient models we observed a decrease in glutarylation and malonylation, which are structurally similar acylations (**Figure 3E**). This data led us to investigate other potential explanations for the acylation patterns observed.

**Figure 3:**
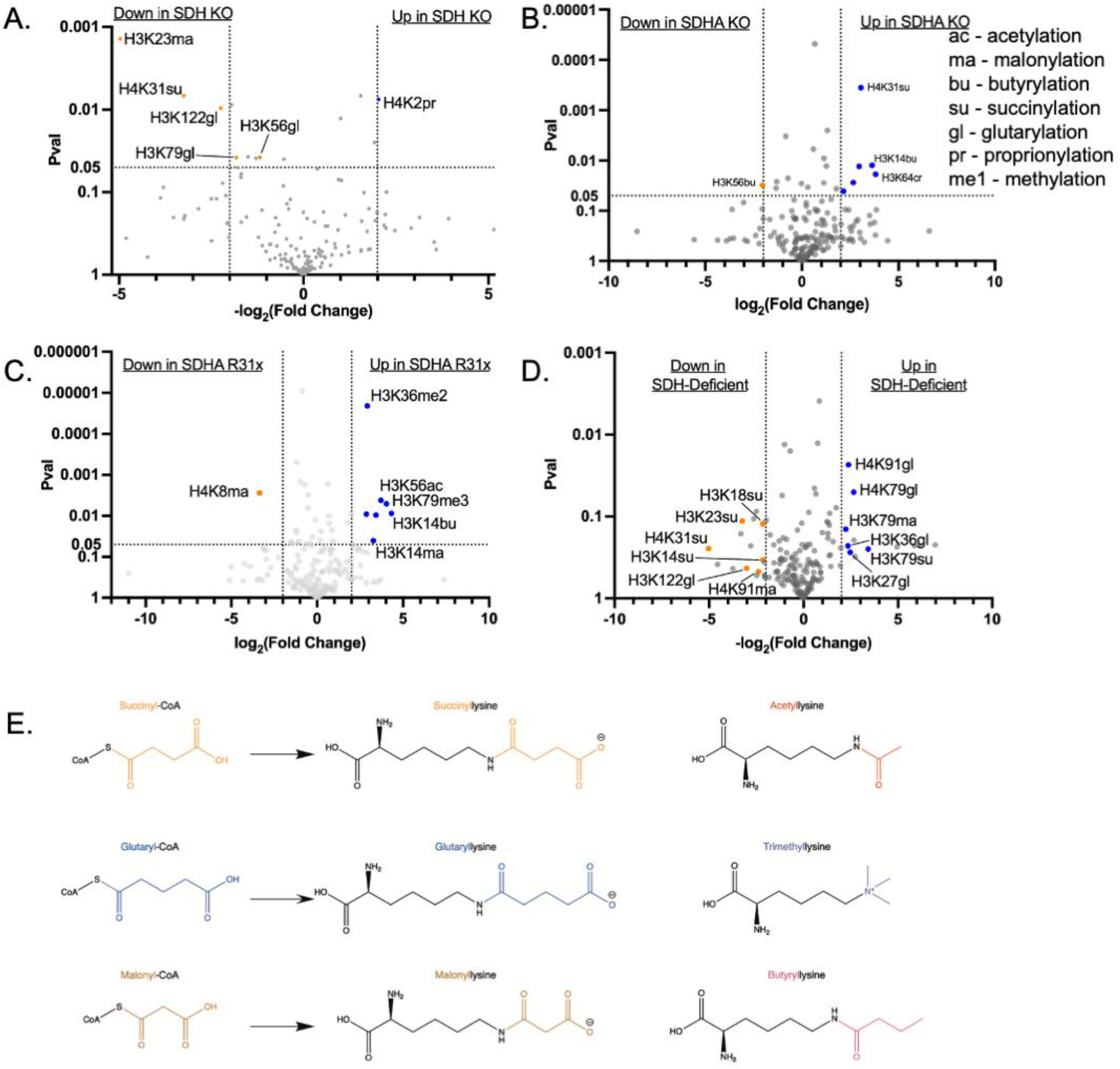
SDH-Deficiency Alters Histone PTM Status Including Decreases In Desuccinylase Sensitive Marks. **A**. Quantification of histone post-translational modification status in 143B SDHB WT and SDHB KO. **B**, in Hap1 SDHA WT and SDHA KO **C**, in Hap1 SDHA WT and SDHA R31x **D**, and in SDH-competent and SDH-deficient GIST patient samples. Desuccinylase sensitive marks, succinylation, glutarylation and malonylation, highlighted. **E**. Chemical structures of free carboxylic short chain acyl-CoAs and their corresponding lysine acylation. P value determined by two-way T-Test for 143B, Hap1, and GIST Patient Histone PTM Analyses. P value < 0.05 and fold change > 4 considered significant.

### SDH-Deficiency Alters Desuccinylase Expression

Given that succinylation can occur both enzymatically and non-enzymatically [15, 31-33], lysine succinylation occurs frequently and is highly sensitive to the concentration of succinyl-CoA [34]. Interestingly, in addition to the observed decrease in histone succinylation, other dicarboxylate histone post translational modifications were decreased, including glutarylation and malonylation. The decrease in histone succinylation status despite the increase in succinyl-CoA led us to hypothesize that major desuccinylases may be upregulated to compensate for the increase of succinyl-CoA. This compensatory upregulation would preserve genome stability and integrity from aberrant succinylation. Previous reports have shown that SIRT7 [35] and class I histone deacetylases (HDAC1-3) [36] are histone desuccinylases, as well as having activity towards dicarboxylate lysine modifications, including malonylation and glutarylation.

We quantified expression of these desuccinylases via western blot and found an upregulation of HDAC2 in the 143B and Hap1 SDH-deficient models (**Figure 4A**). Given that previous work has shown that SIRT7 has desuccinylase activity in the nucleus, we quantified and found no increase in expression in our SDH-deficient 143B and Hap1 models. Additionally, previous work has shown that succinyl-CoA is the predominant acyl-CoA within the mitochondria [37]. Given the increase in succinyl-CoA we observed, there may be off target succinylation that was not captured by our histone PTM proteomic approach. Therefore, we were also interested in desuccinylase expression in the mitochondria. SIRT5 has been reported as a mitochondrial desuccinylase with cross reactivity with glutaryl- and malonyllysine [38]. We found no significant increase in SIRT5 expression in our SDH-deficient 143B and Hap1 cell models (**Figure 4B**). To see if these findings translated to patient samples, we quantified HDACs1/2/3 and SIRT5 and SIRT7 in SDH-competent and SDH-deficient GIST patient samples. Here, we found no significance difference in IHC score between our SDH-competent and SDH-deficient GIST patient samples (**Figure 4C-D**).

**Figure 4:**
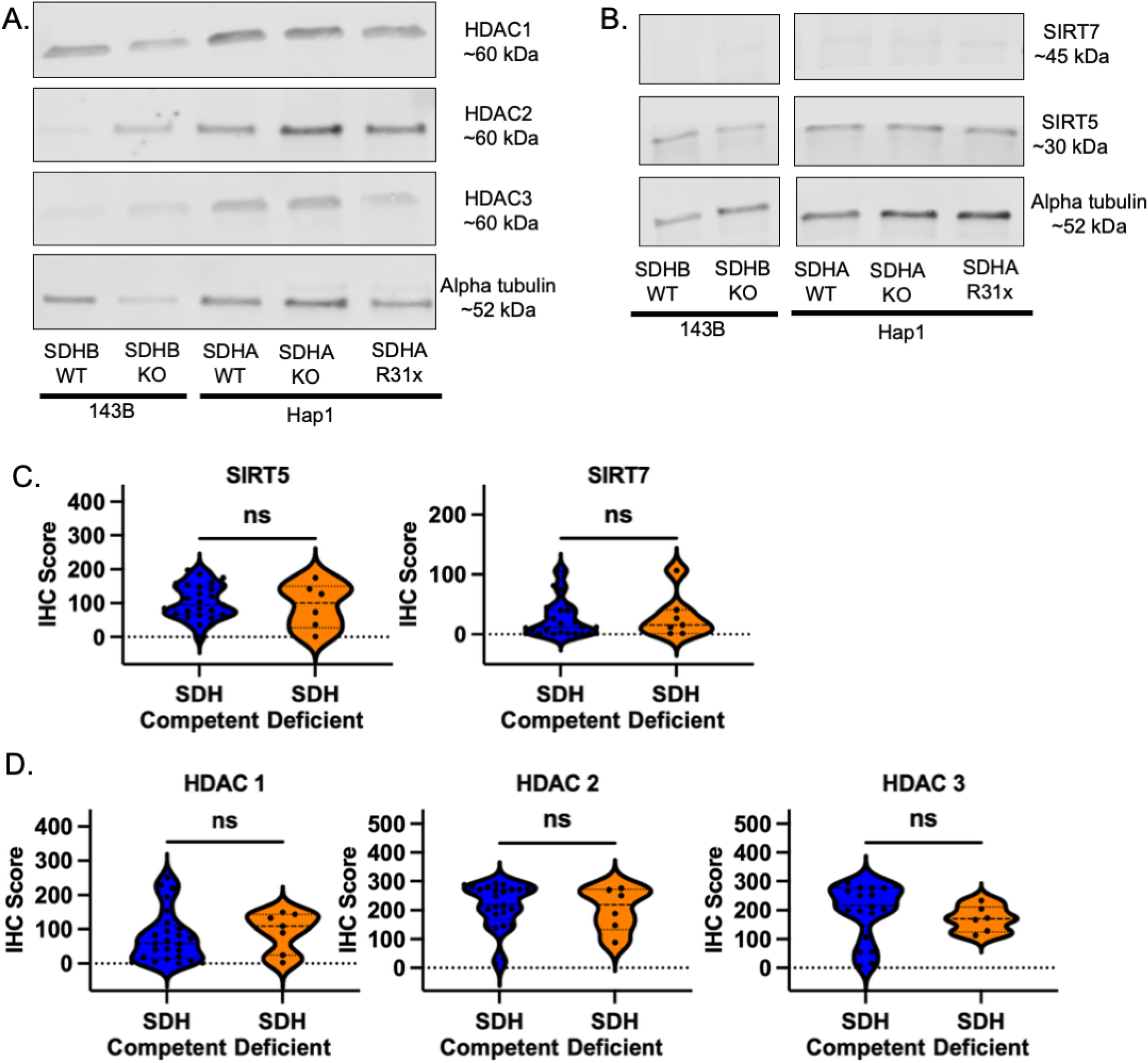
SDH-Deficiency Alters Desuccinylase Expression in 143B and Hap1 SDH-deficient Lines. **A**. Western blot analysis of HDAC 1, HDAC 2, HDAC3 expression in 143B and Hap1 SDH-competent and SDH-deficient cell lines with Alpha-Tubulin as loading control. **B**. Western blot analysis of SIRT7 and SIRT5 in 143B and Hap1 SDH-competent and SDH-deficient cell lines with Alpha-Tubulin as loading control. **C**. Visiopharm IHC score of SIRT5 and SIRT7 in SDH-competent and SDH-deficient GIST patient samples. **D**. Visiopharm IHC score of HDAC1, HDAC 2, HDAC3 in SDH-competent (n = 27) and SDH-deficient GIST patient samples (n = 7). Significance determined by two-way T-Test for GIST Patient IHC Scoring. NS p > 0.05

Taken together, we found that there was an increase in expression of the desuccinylase HDAC2, but not SIRT7 or SIRT5 in our SDH-deficient 143B and Hap1 models. This increase in expression may be a compensatory upregulation of desuccinlyases to prevent aberrant succinylation that may be deleterious to cell viability. As such, despite this, we saw no significant increase in our GIST patient samples, which again may be complicated by tumor heterogeneity.

### SDH-Deficient Cells Are Not Sensitive to HDAC Inhibition

Lastly, we sought to test whether there was an SDH-deficient dependence on desuccinylase activity that may be a therapeutically targetable phenotype. We treated our SDH-competent and SDH-deficient cell model systems with the Class I HDAC inhibitor, trichostatin A (TSA) [39]. We found that 143B and Hap1 SDH-deficient cells had a non-significant decrease in IC50 from 4 nM in the SDH-competent controls to 3-3.5 nM in the SDH-deficient lines (**Figure 5A-B)**. These IC50 values are in line with previous reports in other cell lines [40-42]. We then tested if TSA impacted colony formation in our 143B and Hap1 models. We found no significant difference in colony number, colony area, or area per colony when normalized to vehicle controls, although there were modest decreases in colony number (p=0.09) in the 143B model (**Figure 5C-E**).

**Figure 5:**
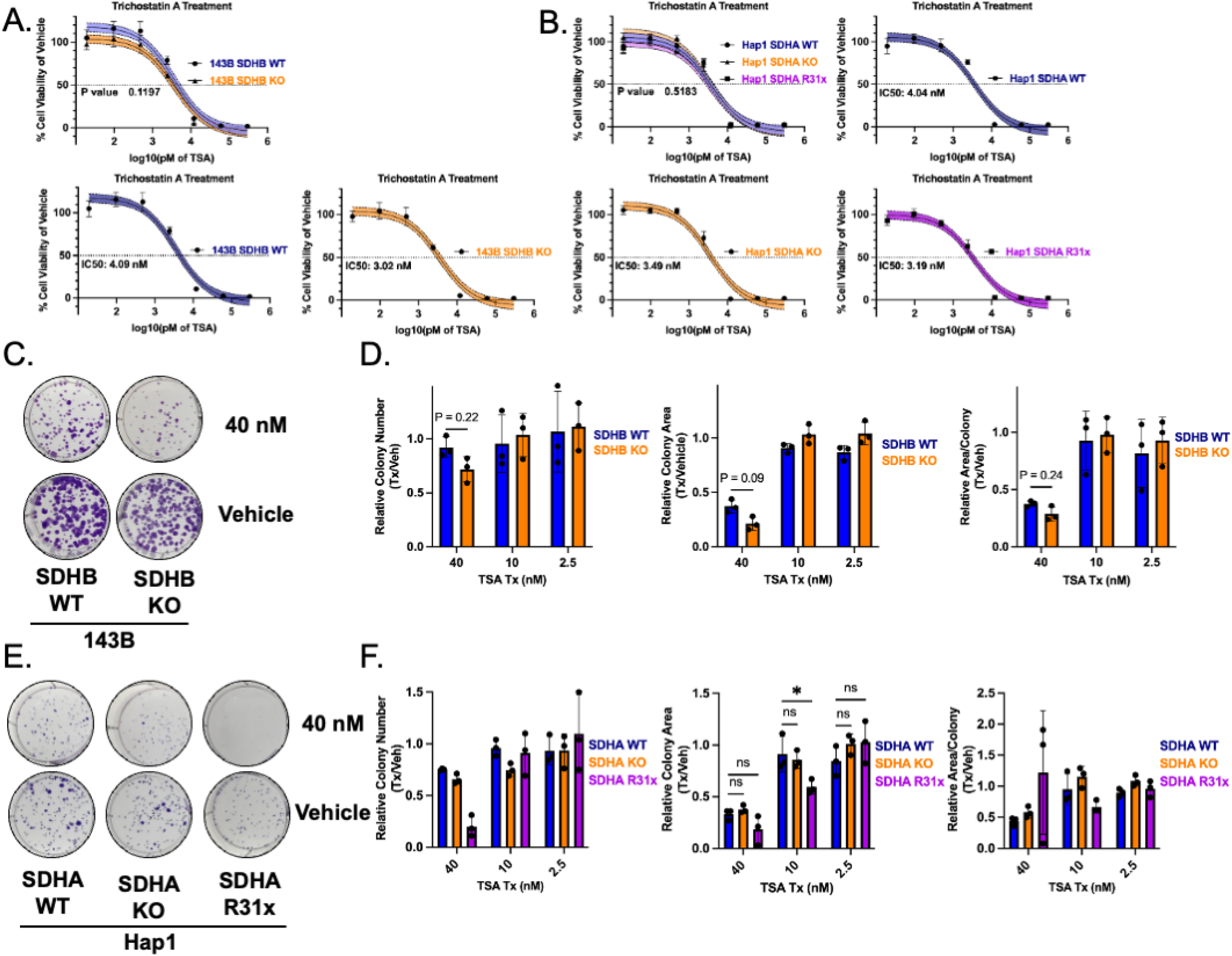
SDH-Deficient Cells Are Not Sensitive to HDAC Inhibition. **A**. Dose response curve of Class I HDAC inhibitor TSA on 143B SDHB WT (n = 6) and SDHB KO (n = 6) cell viability. **B**. Dose response curve of Class I HDAC inhibitor TSA on Hap1 SDHA WT (n = 6), SDHA KO (n = 6), and SDHA R31x (n = 6) cell viability. **C**. Representative images for 143B assay at 40 nM and vehicle (0.1% DMSO) treatment conditions. **D**. Quantification of TSA treatment on colony formation of 143B SDHB WT (n = 3) and SDHB KO (n = 3). **E** and Hap1 formation assay at 40 nM and vehicle (0.1% DMSO) treatment conditions. **F**. Quantification of TSA treatment on colony formation of Hap1 SDHA WT (n = 3) and SDHA KO (n = 3), and SDHA R31x (n = 3). F-test to calculate P-value

## DISCUSSION

In this study we found that succinate dehydrogenase (SDH) deficiency leads to central carbon metabolism rewiring and increases to reactive metabolites, including succinate (**Figure 1A-C**) and succinyl-CoA, the donor for succinylation (**Figure 2A-B**). Additionally, we found no correlation between succinate/fumarate ratio and amount of succinyl-CoA in the NCI-60 cell panel. This may mean that succinate/fumarate ratio is not predictive of succinyl-CoA. Despite our observed succinyl-CoA increase, we found no significant increase in histone succinylation (**Figure 3A-B**). Interestingly, we found a decrease in succinylation and the structurally similar lysine modifications glutarylation and malonylation (**Figure 3E**). Major desuccinylases, including HDAC1/2/3 and SIRT5/7 have been shown to have cross reactivity to glutaryl- and malonyllysines. Due to this, we found an upregulation of HDAC2, but not SIRT5/7 in our 143B and Hap1 SDH-deficient models (**Figure 4A-B**). Additionally, we found that there was no upregulation of desuccinylases in SDH-deficient GIST patient samples when quantified via IHC (**Figure 4C-D**). Lastly, we tested if this upregulation of HDAC2 presented an SDH-deficient therapeutic vulnerability. We quantified the IC50 in our 143B and Hap1 SDH-competent and SDH-deficient model systems following treatment with TSA. We found a modest, but not statistically significant decrease in LD50 (**Figure 5A-B**). Using a colony formation assay, we found that there was again a modest, but not statistically significant, decrease in colony number and colony area in the SDH-deficient 143B and Hap1 models (**Figure 5D-E**). Taken together, these data suggest that there is limited, if any, alterations to SDH-deficient cell viability in response to desuccinylase activity. There may be redundancy in desuccinylase acitivty, so that targeting Class I HDACs alone is not enough to induce SDH-deficient specific cell death. Additionally, there may be unreported desuccinylases that may be able to compensate for loss of Class I HDAC inhibitor in SDH-deficiency. Lastly, there may be additional critical fates for succinyl-CoA beyond histone acylation, including other protein lysine acylations or heme biosynthesis that were unperturbed by tested inhibitors.

Our findings in isogenic cell lines mostly did not align with findings in SDH-deficient GIST patient samples. However, we believe there are potential explanations for the divergence of these findings. Firstly, the heterogeneity of the tumor microenvironment of the SDH-competent and SDH-deficient GIST patient samples may complicate comparisons. We observed an increase in succinate and succinate/fumarate ratio in these patient samples, but not succinyl-CoA. This may be due to the fact that succinate can be exported into the extracellular space and accumulate. Acyl-CoAs are not exported and are rapidly broken down in the extracellular environment [43, 44]. Lastly, previous work has shown that the metabolic and epigenetic consequences of SDH-deficiency may be different depending on the cell context [45]. As such, the immortalized cancer lines 143B and Hap1 may have different metabolic and epigenetic plasticity from the SDH-deficient GIST models. So, while the underlying loss of SDH may be shared, downstream metabolite dependent consequences may be altered.

There are some remaining questions and future directions for this work. Firstly, while we observed an increase in succinyl-CoA on the whole cell level, acyl-CoAs are compartmentalized with specific subcellular compartments having unique acyl-CoA concentrations [46]. As such, our whole cell quantification loses this spatial resolution and there may be compartment specific rewiring of acyl-CoAs. Secondly, we only investigated the histone post-translational modification landscape, but essentially all lysines in the proteome can be acylated [47, 48]. While we observed inconsistent changes in the succinylation status of histones, there may be post-translational modification alterations to other proteins. Lastly, we were not able to spatially map where the post-translational modification alterations are in the genome. Current techniques are reliant on antibody-based assays, which antibodies for specific histone acyllysines have high cross reactivity and low specificities [49-51]. Understanding this spatial distribution would better elucidate gene specific changes to chromatin structure and gene expression.

Taken together, we have identified that our SDH-deficient cells have increased succinyl-CoA concentrations, although this may vary by cell context. Additionally, we found that there is a disconnect between succinyl-CoA concentration and histone succinylation. This disconnect may be driven by an overexpression of desuccinylases, but changes to cell viability upon pharmacological inhibition of these enzymes were modest at best.

## Acknowledgements

NWS was funded by NIH grants R35GM156596, R01CA298386, R01CA292937. LR was funded by R01CA281212. We acknowledge the support of the Metabolomics and Lipidomics Core of the Lewis Katz School of Medicine at Temple University. The Sidoli lab gratefully acknowledges funding from the Aging Biology Foundation, the Hevolution Foundation (AFAR), the ERCM-CFAR Center for AIDS Research, the NIAID for the PROVIDENT grant (1U19AI181977), and the NIH Office of the Director (S10OD03028).

## Author Contributions

A.C., N.W.S., and L.R. conceived the study and wrote the main manuscript. A.C., T.G., S.V., T.C., M.M., J.K., K.H., K.C., S.G., S.St., S.Si., and N.W.S. conducted experiments, performed analysis, and analyzed data. All authors reviewed the manuscript.

